# Near-optimal integration of the magnitude Information of time and numerosity

**DOI:** 10.1101/2023.02.01.526584

**Authors:** Taku Otsuka, Yuko Yotsumoto

## Abstract

Magnitude information is often correlated in the external world, providing complementary information about the environment. As if to reflect this relationship, the perceptions of different magnitudes (e.g., time and numerosity) are known to influence one another. Recent studies suggest that such magnitude interaction is similar to cue integration, such as multisensory integration. Here, we tested whether human observers could integrate the magnitudes of two quantities with distinct physical units (i.e., time and numerosity) as abstract magnitude information. The participants compared the magnitudes of two visual stimuli based on time, numerosity, or both. Consistent with the predictions of the maximum likelihood estimation (MLE) model, the participants integrated time and numerosity in a near-optimal manner; the weights for numerosity increased as the reliability of the numerosity information increased, and the integrated estimate was more reliable than either the time or numerosity estimate. Furthermore, the integration approached a statistical optimum as the temporal discrepancy of the acquisition of each piece of information became smaller. These results suggest that magnitude interaction arises through a similar computational mechanism to cue integration. They are also consistent with the idea that different magnitudes are processed by a generalized magnitude system.

## 1. Introduction

Magnitude information, such as time, space, and numbers, is essential for interacting with the environment. We routinely make conscious and unconscious judgments about how long an event lasted, how large an object is, and how many objects there are. Moreover, different magnitude dimensions are often correlated in the external world and thus provide complementary information about the environment. As if to reflect this relationship in the external world, ample evidence shows that the perception of magnitude in one dimension is influenced by magnitudes in other dimensions. For example, more numerous stimuli are perceived to last longer than less numerous ones, while longer-lasting stimuli are perceived to be more numerous (1–4). In addition to these behavioral interactions between magnitudes, neuroimaging studies have shown that the processing of different magnitudes activates partially overlapping brain regions (5–9). These findings converge into a prevailing theory: “A theory of Magnitude” (ATOM; (10)). ATOM proposes the presence of a common magnitude-processing system that is shared across different dimensions located mainly in the parietal cortex (10,11).

Although ATOM conceptually accounts for the behavioral interactions between magnitudes (often referred to as “magnitude interaction”), its predictions remain ambiguous and therefore cannot explain the systematic differences under different experimental conditions. For example, previous studies suggest that the size and directionality of the magnitude interaction depend on the range of the magnitude (2), the time course of the stimulus (3,12,13), and the reliability of each dimension (14–16). Specifically, the perception of space/numerosity is affected by time when space/numerosity information dynamically accumulates over time (3,12,13,15); the perception of time is influenced by space/numerosity when the reliability (i.e., precision) of space/numerosity is higher than that of time, but it is less affected when the reliability of the two pieces of information is comparable (14–16). Therefore, to explain these observations and further elucidate the mechanism of magnitude interaction, a different framework that enables detailed quantitative predictions is required.

Interestingly, a recent study suggests that magnitude interaction reflects an active “binding” mechanism that integrates different magnitudes of the same stimulus to achieve a unified magnitude representation (17), rather than a contextual interference mechanism as was conventionally assumed (5,10,18,19). This raises the possibility that magnitude interaction (also referred to as “magnitude integration” (3,17)) is considered one form of “cue integration” (20–22). Cue integration typically occurs when a single physical quantity is specified using multiple features that provide redundant information. For example, in a classical “ventriloquist effect,” the perception of the auditory spatial location is dragged by the nearby visual information (23). This effect is thought to reflect the brain’s function of integrating audiovisual information to obtain a single estimate of a spatial location (24,25).

It has been shown that cue integration, including the ventriloquist effect, can be successfully explained by computational models based on a Bayesian framework (24,26–28). A well-known model often tested as a benchmark for cue integration is the maximum likelihood estimation (MLE) model (24,26). Assuming that the estimates from each cue are unbiased and their noises (i.e., variability) are normally distributed and independent of one another, the MLE model makes two important quantitative predictions: First, the integrated estimate (*Ŝ*_12_) is a weighted average of two single estimates (*Ŝ*_1_ and *Ŝ*_2_),

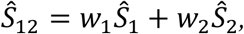

with the weights for cues 1 and 2 (*w*_1_ and *w*_2_) given in proportion to their relative reliabilities:

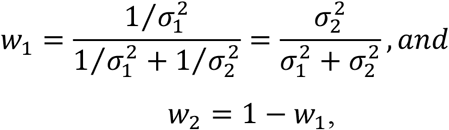

where 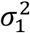 and 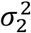 indicate the variances (i.e., the inverse of the reliability) associated with cues 1 and 2, respectively. Second, the variance of the integrated estimate 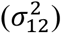 is lower than that of either of the single estimates:

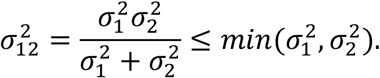

Therefore, the integrated estimate has the lowest possible variance. In this sense, the integration following the MLE model is statistically optimal.

As discussed above, magnitude interaction is qualitatively consistent with the MLE prediction that each piece of information is integrated based on its reliability (14–18). However, no study has directly tested whether the quantitative predictions of the MLE model apply to magnitude integration, where two quantities that have distinct physical units (e.g., time and numerosity) are integrated as abstract magnitude information.

In the present study, we test the hypothesis that the brain integrates the magnitude information of different dimensions according to the MLE model to achieve a unified magnitude representation of the stimulus. We develop a task in which participants are asked to integrate the magnitude information of time and numerosity. In the task, participants compare the magnitude of two visual stimuli based on only time, only numerosity, or both time and numerosity. We manipulate the reliability of the numerosity information by changing the contrast of a set of dots. Moreover, based on a previous report that asymmetries in the time course of a stimulus affect the interaction of time and numerosity (3), we examine the effect of the time course of a stimulus in separate experiments. In Experiment 1, we present the dots (i.e., numerosity) at the onset of the stimulus; in Experiment 2, we present the dots near the offset of the stimulus.

## 2. Methods

### (a) Participants

A total of 42 volunteers (13 females; mean age 20.4, range 18–27 years) participated in the study (21 participants for each of Experiments 1 and 2). All the participants reported normal or corrected-to-normal vision and no history of neurologic or psychiatric conditions. They provided written informed consent before the experiment and received monetary compensation for their participation. The experimental protocol was approved by the Institutional Review Board of the University of Tokyo.

According to a tutorial on cue integration research, 10 participants should be sufficient to test the MLE model (22). However, because previous studies on magnitude integration have tested approximately 15 to 20 participants (2,3,17), we recruited 21 participants for each of Experiments 1 and 2.

### (b) Apparatus and stimuli

The experiments were conducted in a dark room. The stimuli were generated using Psychophysics Toolbox-3 (29) and MATLAB (R2021a; The MathWorks, Inc.) and were presented on a 23.6-inch LCD monitor with 1920 × 1080 resolution at a refresh rate of 120 Hz (VIEWPixx 3D; VPixx Technologies, Inc.). Participants were seated in a chair and their heads fixed on a chin rest 75 cm away from the monitor.

All stimuli were presented on a gray background. Stimuli comprised a set of dots presented on a dynamic random-noise patch. A black fixation cross (0.144 ° radius) was always presented at the center of the screen except during the response period. The dynamic random-noise patch comprised a circle (8.450 ° diameter) and was updated every frame by generating random numbers from a normal distribution. The mean luminance was equal to that of the background, and the contrast was constant under all conditions. Each set of dots contained equal proportions of black and white dots randomly placed in a virtual circle. Each dot had a 0.051 ° radius. The minimum inter-dot and dot-fixation distance was 1.5 times the radius of a single dot, such that each dot did not overlap with any other dot or a fixation cross. The Michelson contrast of the dots was either 100% (high) or 48% (low) in Experiment 1 and either 100% (high) or 55% (low) in Experiment 2 (see “Conditions”). The mean luminance of the dots was equal to that of the background. The radius of the virtual circle on which the dots were presented was varied in each trial to attenuate the correlation between the numerosity and non-numerical properties, such as the convex hull and density; it was either large (4.200 ° radius) or small (3.437 ° radius) for the comparison stimulus, while it was fixed at the average value of large and small for the standard stimulus. An example of stimuli is shown in Figure 1.

**Figure 1.**
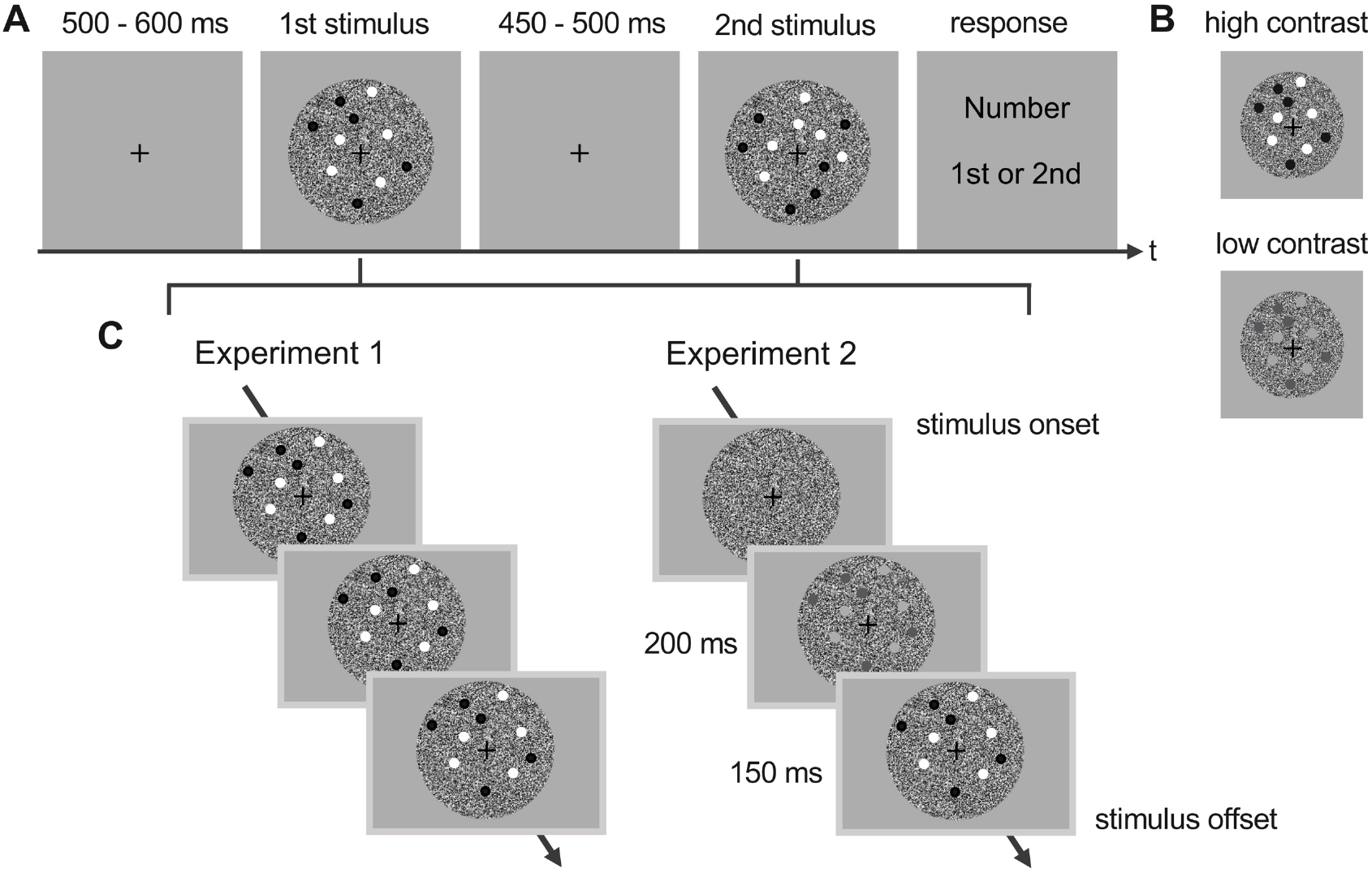
Schematics of the experiments. (A) Trial structure of the task. Participants were sequentially presented with two visual stimuli each comprising a set of dots placed on a dynamic random-noise patch, and were asked to compare the magnitudes of the first and second stimuli based on *time* (i.e., the presentation duration of the dynamic random-noise patch), *numerosity* (i.e., the number of dots), or both *time* and *numerosity*. These three types of trials were presented in separate blocks and participants were informed about what they had to judge before starting each block. The figure depicts an example of a *numerosity* trial. The order of standard and comparison stimuli was counterbalanced. (B) An example of contrast manipulation for numerosity information. The reliability of the numerosity information was manipulated by varying the contrast of the dots in the *numerosity* and *time-numerosity* conditions. The contrast was always high in the *time* conditions. (C) The time course of a stimulus in Experiments 1 and 2. In Experiment 1, a set of dots was presented at the stimulus onset. In Experiment 2, a set of dots was presented 350 ms before the stimulus offset, and its contrast was gradually increased.

### (c) Conditions

Experiments were conducted in a two-interval forced-choice (2IFC) paradigm. In each trial, participants were sequentially presented with two visual stimuli (i.e., standard and comparison) each of which comprised a set of dots placed on a dynamic random-noise patch. Under all conditions, the presentation duration for the dynamic random-noise patch defined the *time* dimension of the stimulus, while the number of dots defined the *numerosity* dimension. Participants were asked to compare the magnitude of the first and second stimuli based on time alone (T), numerosity alone (N), or both time and numerosity (TN). Specifically, in the time conditions (T), participants indicated whether the noise patch of the first or second stimulus lasted longer while ignoring the number of dots. In the numerosity conditions (N), they indicated whether the first or second stimulus comprised more dots while ignoring the duration of the noise patch. In the bidimensional conditions (TN), they attended to both the duration of the noise patch and the number of dots and indicated whether the first or second stimulus lasted longer and comprised more dots. Note that in the TN conditions, the time and numerosity of the comparison stimulus were perfectly correlated (*r* = 1), such that participants could integrate the two dimensions without confusion (see below). Figure 1 shows a visual depiction of a trial.

The standard stimulus lasted 600 ms (*time* dimension) and comprised 24 dots (*numerosity* dimension) in the unidimensional conditions (T, N). In contrast, there were three types of standard stimuli depending on the magnitudes of the *time* and *numerosity* in the TN conditions: Δ = −1, Δ = 0, and Δ = 1, where Δ indicates that *time* was 600 ms multiplied by 1.1^Δ^ while *numerosity* was 24 dots multiplied by 1.1^−Δ^. Therefore, Δ = −1 means that *time* was 600 × 1.1^−1^ ms while *numerosity* was 24 × 1.1^1^ dots; Δ = 1 means that *time* was 600 × 1.1^1^ ms while *numerosity* was 24 × 1.1^−1^ dots; and Δ = 0 means that *time* was 600 ms while *numerosity* was 24 dots. The comparison stimulus took nine possible magnitude values, defined as the 600 ms and 24 dots multiplied by 1.1^−5^, 1.1^−3^, 1.1^−2^, 1.1^−1^, 1.1^0^, 1.1^1^, 1.1^2^, 1.1^3^, and 1.1^5^, resulting in durations of 375, 450, 500, 542, 600, 658, 725, 800, and 966 ms (*time* dimension) and numerosities of 15, 18, 20, 22, 24, 26, 29, 32, and 39 dots (*numerosity* dimension). This manipulation was necessary to make the magnitudes of *time* and *numerosity* comparable on a logarithmic scale (these values were rounded off to the nearest integer and the durations corresponded to the presented durations calculated from the monitor’s refresh rate). In the T conditions, only the *time* dimension of the comparison stimulus could vary while *numerosity* was fixed at the standard magnitude (24 dots); in the N conditions, only *numerosity* could vary while *time* was fixed at the standard magnitude (600 ms). In the TN conditions, both *time* and *numerosity* could vary, but they were perfectly correlated (*r* = 1). That is, *time* and *numerosity* were identical in magnitude relative to the standard magnitude.

To manipulate the reliability of the numerosity information, we used two levels of dot contrast, high and low, in the N and TN conditions (see “Apparatus and stimuli” and Figure 1B). The dot contrast was fixed at the high value in the T conditions. Within trials, standard and comparison stimuli had the same contrast.

To examine the effect of the time course of a stimulus on magnitude integration, we conducted two experiments. In Experiment 1, a set of dots was always presented from the stimulus onset to the stimulus offset. In Experiment 2, a set of dots was presented 350 ms before the stimulus offset, gradually increasing its contrast over 200 ms, reaching the specified contrast (i.e., high or low) 150 ms before the stimulus offset (see Figure 1C). Experiments 1 and 2 differed in terms of how much of a gap there was between when *time* and *numerosity* information became available; *time* was available only at the offset, *numerosity* was always available from the onset in Experiment 1, while *numerosity* also became available near the offset in Experiment 2. In other words, the gap in the timing for when *time* and *numerosity* information became available was smaller in Experiment 2 than in Experiment 1.

### (d) Procedure

The conditions (T, N, and TN) were presented in separate blocks. Before each block, participants were cued on whether they had to make judgments about *time*, *numerosity*, or both *time* and *numerosity*. In each trial, the standard and comparison stimuli were presented with an inter-stimulus interval (ISI) sampled from a uniform distribution between 450 and 550 ms. The order of the standard and comparison stimuli was counterbalanced across trials. Participants were instructed to press *M* on a conventional keyboard if they perceived the second stimulus as larger in magnitude than the first stimulus, and *C* if they perceived the first stimulus as larger in magnitude than the second stimulus. After the response, the next trial started with an inter-trial interval sampled from a uniform distribution between 500 and 600 ms. In the actual experiment, no feedback was provided regarding the correct or incorrect responses.

Each experiment (i.e., Experiments 1 and 2) was conducted in two sessions on two different days. Each session comprised 21 blocks. These 21 blocks were further divided into three sub-blocks each of which comprised one block of T trials, two blocks of N trials, and four blocks of TN trials. The order of the blocks within the sub-blocks was randomized. Each block comprised either 36 trials (T and N blocks) or 54 trials (TN blocks). The two levels of dot contrast (high and low) and three types of standard stimuli (Δ = −1, Δ = 0, and Δ = 1) were randomly presented within the same block. Specifically, each T block comprised 9 (comparison magnitude) × 4 (repetition) trials, each N block comprised 9 (comparison magnitude) × 2 (dot contrast, high and low) × 2 (repetition) trials, and each TN block comprised 9 (comparison magnitude) × 2 (dot contrast, high and low) × 3 (standard stimuli, Δ = −1, Δ = 0, and Δ = 1) trials. Thus, one data point of a psychometric function was calculated from a proportion of 24 trials. In total, 1,944 trials were performed per participant.

To familiarize the participants with the task and to allow them to learn that *time* and *numerosity* were perfectly correlated in the TN condition, each experimental session for both Experiments 1 and 2 was preceded by a short practice session. The practice session comprised four blocks of T, N, and two TN conditions with the standard stimulus of Δ = 0. After every response, the participants received feedback, i.e., the fixation cross was changed to green/red for 200 ms to indicate a correct/incorrect response.

### (e) Data analysis

For each participant and every 18 conditions, the proportion of trials for which the comparison stimulus was judged as larger was calculated as a function of the comparison magnitude. These data were fitted to a cumulative normal distribution to obtain a psychometric function using Palamedes toolbox 1.10.11 (30). From the fitted psychometric functions, the point of subjective equality (PSE) and just-noticeable difference (JND) were obtained. The PSE corresponded to the magnitude at which the probability of “larger” responses reached 50%. The JND was defined as the inverse of the slope parameter (*JND* = 1⁄*β*) of the fitted function. The empirical variance, 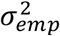, was then computed using the following formula for each condition (22):

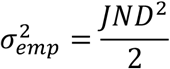

In the main analysis, we used the sensory variances obtained from the unidimensional conditions (T and N) to determine the predicted weights and variances under the bidimensional conditions (TN). The predicted numerosity weights, *W_N,pred_*, were computed as follows:

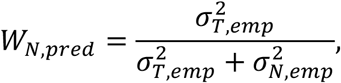

where 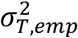 and 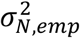 were empirical variances under the T and N conditions, respectively. The predicted bidimensional variances, 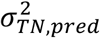, were computed according to the following formula:

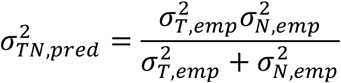

The empirical numerosity weights were calculated from the bidimensional conditions according to the method used in (31). First, PSEs were measured for the standard stimuli of Δ = −1, Δ = 0, and Δ = 1, and were plotted as a function of Δ. A regression line was then fitted to the data, and the empirical numerosity weights, *W_N,emp_*, were computed as follows:

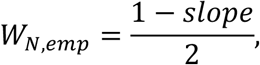

where *slope* was the estimated slope of the regression line (31). The empirical variance in the TN conditions, 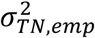, was defined as the average of the variances for Δ = −1, Δ = 0, and Δ = 1 (24). Note that in the subsequent analysis, the standard deviation (*σ*) rather than the variance was used as an index of the variability parameter.

For statistical testing, a two-way repeated-measures ANOVA was conducted on the weights and variability, with prediction (predicted and empirical) and contrast (high and low) as within-subjects factors. For each dot-contrast level, the predicted and empirical weights/variability were compared using two-sided paired-sample t-tests. Moreover, for each dot-contrast level, the empirical variability in the TN conditions (*σ_TN,emp_*) was compared with the minimum value of the unidimensional variability (*σ_T,emp_*), (*σ_N,emp_*)) using two-sided paired-sample t-tests. Additionally, we calculated the correlation coefficients between the predicted and empirical weights/variability. These statistical tests were further assessed by evaluating the null hypothesis and the strength of the evidence by computing Bayes factors using JASP (32). BF_10_ represents evidence in favor of the alternative hypothesis that there is a difference/correlation. BF_10_ > 3 indicates support for the alternative hypothesis, whereas BF_10_ < 1/3 indicates support for the null hypothesis that there is no difference/correlation.

## 3. Results

### (a) Experiment 1

Figure 2A shows the across-participants mean of the psychometric function in the unidimensional (T, N) and bidimensional (TN) conditions. As expected, the slope of the psychometric function was flatter when the dot contrast was low than when it was high, indicating that the reliability of the numerosity information varied with the dot contrast. Based on the PSEs and JNDs obtained from these psychometric functions, we computed the predicted weights and variability according to the MLE model and compared them with the empirical values (see “Data analysis”).

**Figure 2.**
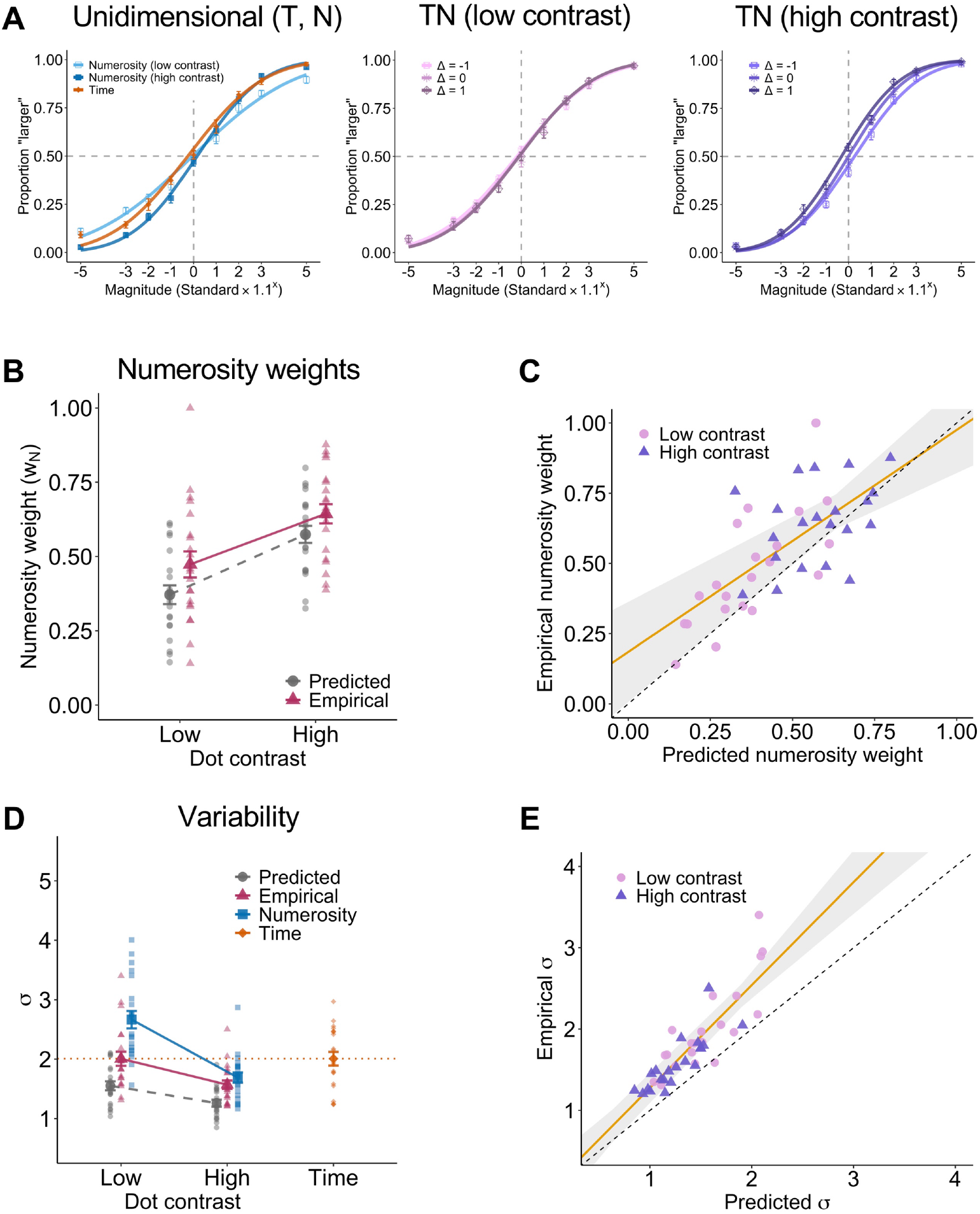
Psychometric functions, numerosity weights, and variability in Experiment 1. (A) The across-participants mean of the psychometric function in the unidimensional (time, numerosity) and bidimensional (time–numerosity) conditions separately for each contrast level and Δ. The Δ in the bidimensional conditions indicates that *time* for the standard stimulus was 600 ms multiplied by 1.1^Δ^ while *numerosity* was 24 dots multiplied by 1.1^−Δ^ (see the “Methods” section). The proportion of trials for which the comparison stimulus was judged as larger in magnitude was plotted as a function of the comparison magnitude. (B) The across-participants mean of the predicted (circles) and empirical (triangles) numerosity weights. Each data point represents an individual participant. (C) Individual predicted and empirical numerosity weights. Symbols indicate contrast levels: high (triangles) and low (circles). The solid line indicates the linear fit of the empirical weights to the predicted weights. The shaded region indicates the 95% confidence interval for the linear fit. (D) The across-participants mean of the predicted variability (circles) and empirical variability in the bidimensional (triangles), time (diamonds), and numerosity (squares) conditions. Each data point represents an individual participant. (E) Individual predicted and empirical variability. Symbols indicate contrast levels: high (triangles) and low (circles). The solid line indicates the linear fit of the empirical variability to the predicted variability. The shaded region indicates the 95% confidence interval for the linear fit. Error bars indicate the standard error of the mean (SEM).

#### Weights

Figure 2B shows the predicted and empirical numerosity weights. A repeated-measures ANOVA revealed significant main effects for prediction and for contrast: *F*(1, 20) = 13.28, *p* = .002, BF_10_ = 71.77, 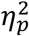 = 0.399; and *F*(1, 20) = 72.99, *p* < .001, BF_10_ > 10000, 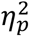 = 0.785, respectively. There was no significant interaction: *F*(1, 20) = 0.61, *p* = .445, BF_10_ = 1.48, 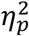 = 0.030. As expected, numerosity weights were larger when the contrast was higher. However, the empirical numerosity weights were larger than the predicted weights when the contrast was low: *t*(20) = 3.39, *p* = .003, BF_10_ = 14.37, *d* = 0.74; however, they were not significantly different when the contrast was high: *t*(20) = 2.03, *p* = .056, BF_10_ = 1.25, *d* = 0.44. Moreover, there was a significant correlation between the predicted and empirical numerosity weights: *r* = 0.692, *p* < .001, BF_10_ > 10000 (Figure 2C).

#### Variability

Figure 2D shows the predicted and empirical variability (σ) in the bidimensional conditions as well as the variability in the unidimensional conditions. A repeated-measures ANOVA revealed significant main effects for prediction and for contrast: *F*(1, 20) = 63.26, *p* < .001, BF_10_ > 10000, 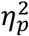 = 0.760; and *F*(1, 20) = 49.99, *p* < .001, BF_10_ > 10000, 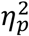 = 0.714, respectively, and a significant interaction: *F*(1, 20) = 6.00, *p* = .024, BF_10_ = 3.40, 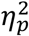 = 0.231). As expected, the variability in the N and TN conditions was smaller when the contrast was higher. However, the empirical variability was systematically larger than predicted. The deviation was more profound when the contrast was high than when it was low: *t*(20) = 7.73, *p* < .001, BF_10_ > 10000, *d* = 1.69 (high contrast), and *t*(20) = 6.58, *p* < .001, BF_10_ = 9258.65, *d* = 1.44 (low contrast).

Moreover, the bidimensional variability was not smaller than the smallest variability in the unidimensional conditions; when the contrast was low, the bidimensional variability was not significantly different from the variability of time alone: *t*(20) = 0.01, *p* = .996, BF_10_ = 0.23, *d* = 0.00; and when the contrast was high, it was not significantly different from the variability of numerosity alone: *t*(20) = 2.03, *p* = .055, BF_10_ = 1.26, *d* = 0.44. Nevertheless, there was a significant correlation between the predicted and empirical variabilities: *r* = 0.862, *p* < .001, BF_10_ > 10000 (Figure 2E).

These results suggest that the MLE predictions are partially supported for both weights and variability. However, the numerosity weights and bidimensional variability were systematically larger than predicted. Furthermore, we found no evidence to support a variance reduction due to integration, which was the critical prediction of the MLE model.

### (b) Experiment2

Figure 3A illustrates the across-participants average of the psychometric function. As in Experiment 1, the psychometric function indicated that the reliability of the numerosity information varied with the dot contrast.

**Figure 3.**
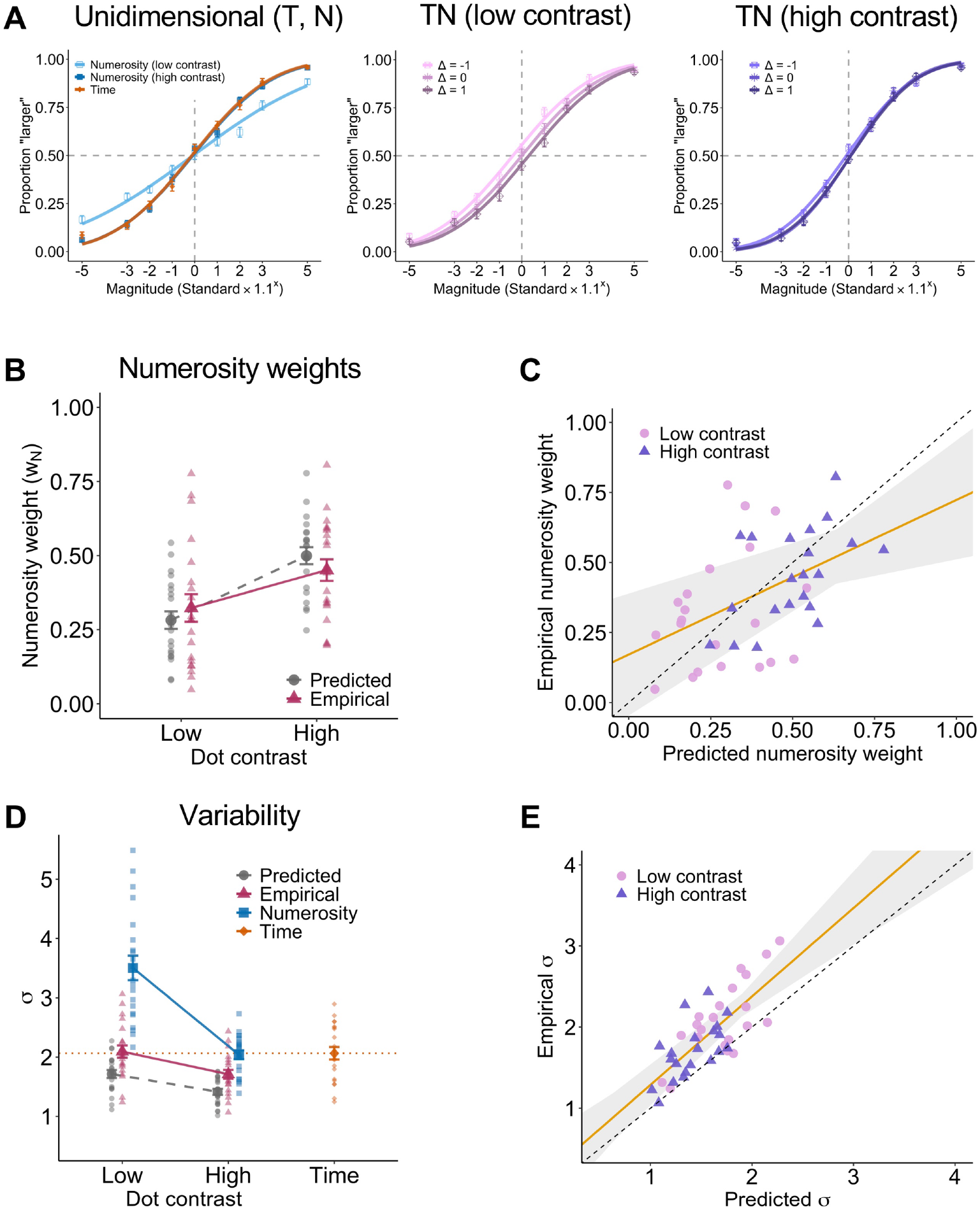
Psychometric functions, numerosity weights, and variability in Experiment 2. (A) The across-participants mean of the psychometric function in the unidimensional and bidimensional conditions separately for each contrast level and Δ. (B) The across-participants mean of the predicted (circles) and empirical (triangles) numerosity weights. Each data point represents an individual participant. (C) Individual predicted and empirical numerosity weights. Symbols indicate contrast levels: high (triangles) and low (circles). The solid line indicates the linear fit of the empirical to the predicted weights. The shaded region indicates the 95% confidence interval for the linear fit. (D) The across-participants mean of the predicted variability (circles) and empirical variability in the bidimensional (triangles), time (diamonds), and numerosity (squares) conditions. Each data point represents an individual participant. (E) Individual predicted and empirical variabilities. Symbols indicate contrast levels: high (triangles) and low (circles). The solid line indicates the linear fit of the empirical to the predicted variability. The shaded region indicates the 95% confidence interval for the linear fit. Error bars indicate SEM.

#### Weights

Figure 3B shows the predicted and empirical numerosity weights. A repeated-measures ANOVA revealed a significant main effect for contrast: *F*(1, 20) = 46.20, *p* < .001, BF_10_ > 10000, 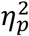 = 0.698. In support of the MLE model, there was no main effect for prediction: *F*(1, 20) = 0.01, *p* = .918, BF_10_ = 0.27, 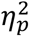 = 0.001; neither was there a main effect for interaction: *F*(1, 20) = 3.65, *p* = .071, BF_10_ = 0.59, 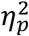 = 0.154. Confirming these observations, we found no evidence of a difference between the empirical and predicted numerosity weights: *t*(20) = 1.49, *p* = .152, BF_10_ = 0.59, *d* = 0.36 (high contrast) and *t*(20) = 0.85, *p* = .403, BF_10_ = 0.32, *d* = 0.19 (low contrast). Moreover, there was a significant correlation between the predicted and empirical numerosity weights: *r* = 0.476, *p* = .001, BF_10_ = 25.71 (Figure 3C).

#### Variability

Figure 3D shows the predicted and empirical variabilities (σ). A repeated-measures ANOVA revealed significant main effects for prediction and for contrast: *F*(1, 20) = 34.04, *p* < .001, BF_10_ > 10000, 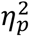 = 0.630; and *F*(1, 20) = 66.53, *p* < .001, BF_10_ > 10000, 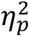 = 0.769, respectively. There was no significant interaction: *F*(1, 20) = 1.94, *p* = .179, BF_10_ = 1.69, 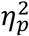 = 0.088). As in Experiment 1, the empirical variability was systematically larger than the predicted variability: *t*(20) = 4.98, *p* < .001, BF_10_ = 366.72, *d* = 1.09 (high contrast) and *t*(20) = 5.41, *p* < .001, BF_10_ = 891.48, *d* = 1.18 (low contrast). Importantly, the bidimensional variability was smaller than either of the unidimensional variabilities when the contrast was high: *t*(20) = 3.82, *p* = .001, BF_10_ = 33.70, *d* = 0.83 (vs. numerosity) and *t*(20) = 3.69, *p* = .001, BF_10_ = 26.27, *d* = 0.81 (vs. time); however, it was not different from the variability of time alone when the contrast was low: *t*(20) = 0.30, *p* = .769, BF_10_ = 0.24, *d* = 0.07. Moreover, there was a significant correlation between the predicted and empirical variabilities: *r* = 0.753, *p* < .001, BF_10_ > 10000 (Figure 3E).

In summary, the results demonstrated that the empirical weights were consistent with the predictions. However, the empirical variability was systematically larger than the predicted variability. Most importantly, the variability in the bidimensional condition was smaller than that in the time and numerosity conditions, supporting the MLE prediction of a variance reduction due to integration.

## 4. Discussion

This study aimed to test whether the integration of different magnitude information was statistically optimal, as predicted by the MLE model. As a test case, we examined the integration of time and numerosity. We found the following results from two experiments: First, when there was a large discrepancy in the timing of the acquisition of the time and numerosit*y* information, numerosity was afforded more weight than predicted; the bidimensional variability was higher than predicted and comparable to the most reliable unidimensional variability. Second, as the discrepancy in the timing became smaller, the numerosity weights approached the MLE predictions; the bidimensional variability remained higher than predicted, yet smaller than the most reliable unidimensional variability.

These results suggest that human observers can integrate different magnitude information in a near-optimal manner. In previous studies, the MLE model has accounted for the integration of multiple cues that provide redundant information about a single physical quantity, such as the visual-haptic integration of length (26), audiovisual integration of spatial location (24), and integration of various visual cues to depth (33,34). In contrast, the current study has demonstrated that two quantities with different physical units, time and numerosity, are near-optimally integrated as abstract magnitude information. Furthermore, the differences between the results of our two experiments suggest that the coincidence of the timing of the acquisition of each piece of information is crucial for optimal magnitude integration. Interestingly, spatial-temporal discrepancies in the acquisition of information have been shown to affect optimal integration, even in the multisensory integration of single physical quantities such as spatial location and orientation (35–37). For example, Plaisier et al. (37) reported that, in the visual-haptic integration of surface orientation, optimal integration breaks down when there is a discrepancy between the visual and haptic exploration modes (i.e., the instantaneous perception of the surface orientation by touching/seeing two spots vs. sequential perception by tracing the surface). This suggests that the time-numerosity interaction may result from a similar computational mechanism to the multisensory integration of single physical quantities.

In contrast to the multisensory integration of a single physical quantity, one might consider the integration of different magnitude dimensions to be outside the scope of the application of the MLE model. However, even in the audiovisual integration of spatial location, visual and auditory signals are assumed to be transformed into a common body-coordinate system (21). This is conceptually similar to assuming that time and numerosity are transformed into a common magnitude representation (10). Therefore, we believe it would be reasonable to apply the MLE model to the integration of multiple magnitude dimensions that have distinct units.

However, it should be noted that the near-optimal integration observed in our study does not necessarily mean that the magnitude integration shares a common mechanism with the cue integration of other features. In particular, the present study cannot determine whether the magnitude integration occurs at the perceptual or at the post-perceptual decision level. However, our results indicate that the magnitude integration is affected by the timing at which information is acquired, suggesting that it could not be explained solely by the decision-level mechanisms, such as pure cognitive reasoning. Moreover, considering that magnitude interaction occurs not only at the decision level but also at the perceptual level (38,39), it is possible that magnitude integration partly involves perception-level mechanisms.

In addition to the MLE model, our findings are consistent with those obtained in previous studies on magnitude interaction. Separate studies have hypothesized that the direction and size of magnitude interaction are affected by the processing time course (3,12) and reliability (14,15,40) of each piece of information. However, the processing time course and reliability have not been independently manipulated within a single study, leaving uncertain the contribution of each factor to the interaction. For example, Togoli et al. (3) reported that a bidirectional interaction between time and numerosity occurred when numerosity information dynamically unfolded over time. While this result suggests that the time course of the stimulus and neural processing dynamics it entails are important factors, dynamically unfolding numerosity information was accompanied by reduced reliability, and thus the results could also be explained by the reliability of the numerosity information. Meanwhile, Cai and Wang (15) proposed that the crucial factor for magnitude interaction was the representational noise (i.e., reliability). However, their experiments used dynamic stimuli to manipulate reliability, confounded by the time course of the processing of the stimulus. In contrast, our study independently manipulated the reliability and time course of the stimulus by varying the contrast of the dots and timing at which numerosity information became available. We found that numerosity weights flexibly varied with reliability (i.e., dot contrast), consistent with studies claiming the influence of reliability on magnitude interaction (14,15,40). Moreover, integration was affected by the timing at which each piece of information was acquired, showing the influence of the time course of the processing of the stimulus. In addition, our results for the near-optimal integration provide further support for the recent hypothesis that magnitude interaction reflects active magnitude binding rather than passive contextual interference between dimensions (17).

Overall, the results of both the present and previous studies suggest that magnitude interaction is caused by an adaptive mechanism that flexibly changes with the reliability and time course of a stimulus. Furthermore, it may reflect an integration mechanism to obtain a unified representation of the stimulus magnitude (17) rather than an interference effect (16,18). However, a caveat to our study is that we used a different experimental paradigm from the one often used to study magnitude interaction. In most previous studies, participants compared the magnitude of one specific dimension while the other dimensions independently varied. In the present study, however, the magnitudes of the two dimensions (i.e., time and numerosity) were perfectly correlated, and the participants were explicitly asked to integrate them. While these manipulations were necessary as they were prerequisites for the MLE model, they were quite artificial compared to the relationship between the magnitudes in the real world; in reality, different magnitudes do not necessarily correlate with one another, and there are situations in which they should not be integrated. In this sense, the generalizability of the present results is limited to the situation in which the two magnitudes must be integrated. However, studies on multisensory integration have developed Bayesian models that explain adaptive behavior in situations in which two signals should not necessarily be integrated (25,27,28,41). Therefore, similar Bayesian models could also describe magnitude interaction (c.f., Cai et al., 2018; Chen et al., 2016, 2021). Regarding the existing theory, the present results support the basic idea of ATOM (10) that different magnitude information is integrated into a common representation, which subsequently contributes to efficient motor output (11). However, the MLE model assumes independence of the noise associated with each piece of information, whereas ATOM suggests that the encoding mechanism is partially shared across different magnitude dimensions (10,11). As it is unlikely that these two assumptions can be simultaneously satisfied, further theoretical development may require a modification of either model.

Incidentally, in the present study, the empirical variability was consistently higher than that predicted by the MLE, although there was a variance reduction apparently due to integration. This suggests that magnitude integration may not satisfy some of the MLE assumptions (22). Potential reasons include the possibilities that (1) there was a mix of subjects/trials that properly integrated the two magnitudes and subjects/trials that relied only on the reliable dimension without integrating the two magnitudes, (2) the MLE model assumption that noises are normally distributed does not hold true for time and numerosity representations, and (3) there is a correlation between the noises associated with the time and numerosity representations. In particular, the third possibility seems consistent with the prediction of ATOM that there is a partially common encoding mechanism for time and numerosity. Future research should consider each of these possibilities and further elaborate on the relationship between the MLE model and ATOM.

## Ethics

The experimental protocol was approved by the Institutional Review Board of the University of Tokyo (#258-8) and was in accordance with the Declaration of Helsinki. All the participants provided written informed consent before the experiment.

## Data accessibility

The data, scripts, and details of the statistical results are available on Open Science Framework: https://osf.io/bx4qt/ (44)

## Acknowledgements

We would like to thank Editage (http://www.editage.com) for English language editing.

